# Direct Visualization of Protein Kinase A Activation on DNA Origami using Fast-Scan Atomic Force Microscopy

**DOI:** 10.1101/2020.09.07.286070

**Authors:** Ioanna Mela, Daniel Ladant, Hiroshi Sugiyama, Masayuki Endo, J. Michael Edwardson, Robert M. Henderson

**Affiliations:** Department of Pharmacology, University of Cambridge, Cambridge CB2 1PD, United Kingdom; Unité de Biochimie des Interactions Macromoléculaires, Département de Biologie Structurale et Chimie, Institut Pasteur, CNRS, UMR 3528, 28 Rue du Dr. Roux, Paris, 75015, France; Department of Chemistry, Graduate School of Science, Kyoto University, Kitashirakawa-oiwakecho, Sakyo-ku, Kyoto 606-8502, Japan; Institute for Integrated Cell Material Sciences, Kyoto University, Yoshida-ushinomiyacho, Sakyo-ku, Kyoto, 606-8501, Japan

**Keywords:** Atomic force microscopy, Fast Scan AFM, PKA, signalling

## Abstract

DNA origami nanostructures provide a unique platform for the direct observation of protein-protein interactions at the single-molecule level. Here, we used DNA origami in combination with fast-scan atomic force microscopy to observe the activation-induced dissociation of individual protein kinase A (PKA) holoenzymes. The PKA holoenzyme consists of two regulatory (R) and two catalytic (C) subunits. When cAMP binds to the R subunits it causes dissociation of the C subunits from the R subunit dimer and activation of the enzyme. Using a DNA origami platform, we were able to observe the activation of PKA in response to photolysis of caged cAMP. Furthermore, exploiting the potential of DNA origami for precise positioning of biomolecules, we were able to position the catalytic subunit of adenylyl cyclase in close proximity to PKA and to observe the activation of PKA in response to cAMP produced by adenylyl cyclase. We provide tools for the observation of signalling pathways at the single-molecule level and show that the C subunits of PKA dissociate from the holoenzyme but stay within ~10 nm of the R subunit dimer upon activation.

---

The great potential of DNA origami nanostructures for functionalization, and the variety of attachment mechanisms available^1–8^, have given rise to a number of origami-based studies of single-molecule interactions *in vitro*^7, 9–11^. Here, we exploit the precise control of protein positioning on DNA origami tiles to observe the activation of protein kinase A (PKA) by cAMP at the single-molecule level. PKA is one of the simplest kinases, and so has served as a model for the study of other members of the protein kinase family^12^. It is a key component in intracellular signalling and in the control of metabolism, and dysregulation of PKA is thought to play a role in various diseases^13^, and in ageing^14^.

The PKA holoenzyme consists of two regulatory (R) and two catalytic (C) subunits^15^. The R subunits serve two main roles: capture of the C subunits and consequent inhibition of their catalytic activity, and localization of the PKA holoenzyme through binding to AKAP scaffold proteins^16–19^. Each of the R subunits contains two cAMP binding sites. Once four cAMP molecules are bound to the R subunit dimer, the two C subunits become activated^20^. Although PKA is a very well-studied protein, there are still unanswered questions about its mechanism of action, in particular regarding the dynamics of the interaction between the C subunits and the R subunit dimer during the activation/deactivation cycle. Two recent studies on the potential mechanisms underlying this process in the cellular context have generated conflicting results^21,22^. Specifically, Smith *et al*.^21^ reported that at physiological concentrations of cAMP (1-2 μM) 70-80% of the C subunits remain associated with the AKAP/R subunit complex, and that substantial release of the C subunit was only seen at supraphysiological concentrations of cAMP (10-90 μM). In contrast, Walker-Gray *et al*.^22^ found that the C subunits are rapidly released upon activation and then recaptured.

Here, we use single-molecule imaging by fast-scan atomic force microscopy (FS-AFM) to demonstrate that 2-D DNA nanostructures can be used as platforms on which to observe signalling at the single-molecule level. We use this technology to observe the behavior of the C subunits of PKA during activation, in near-real time.

We designed and synthesized a rectangular (100 × 70 nm) DNA origami ‘tile’ onto which we could place the catalytic domain of adenylyl cyclase and PKA at a distance of ~20 nm apart (Figure 1.A). The tile carries an ‘orientation loop’ on one side so that the attachment of proteins to the target region can be assessed (Figure 1.B). Tile assembly occurred with an efficiency of 92±5% (n=5 experiments, 100 tiles each, Supplementary Figure 1). The two proteins of interest are attached onto their specific positions on the origami tile through streptavidin-biotin interactions for adenylyl cyclase and a SNAP tag-benzylguanine interaction for PKA (Figure 1.C).

**Figure 1.**
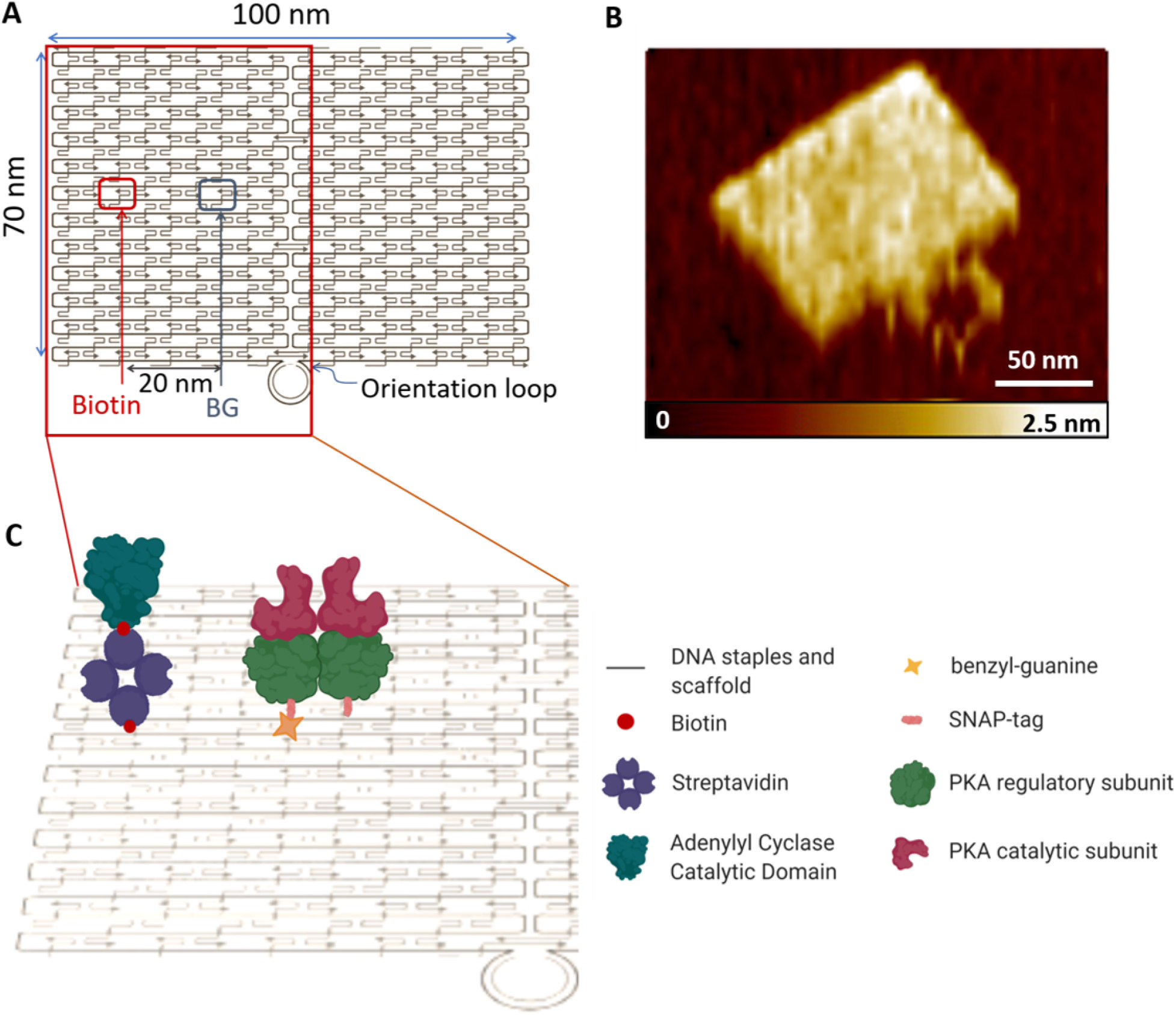
DNA origami template and protein modification. **1.A:** Map of the DNA origami tile (100 × 70 nm) used in the present study. The tile carries biotin and benzylguanine (BG) modifications, ~20 nm apart, for the attachment of adenylyl cyclase and PKA, respectively. **1.B:** Representative AFM image of an origami tile showing the orientation loop. **1.C:** Schematic representation of the protein assembly used for this study. The biotinylated catalytic domain of adenylyl cyclase is attached to the origami tile through a biotin-streptavidin bridge while the PKA holoenzyme is held on the tile through a SNAP tag modification of the R subunit, which binds a benzylguanine-modified staple. Created with BioRender.com

We used a stepwise approach to protein attachment. Following incubation of the tiles with streptavidin, we found that 96±8% (5 experiments, 250 tiles in total) of the tiles carried a streptavidin molecule, positioned at the known location of the biotinylated staple, on the edge of the tile, to the left of the orientation loop (Figure 2.A). The high efficiency of streptavidin binding was expected, given the high affinity of streptavidin for biotinylated targets^23–25^. The peak volume of the bound molecules was 124±3 nm^3^ (n=80), close to the calculated theoretical volume of a streptavidin tetramer (molecular mass, 55 kDa; theoretical volume, 105 nm^3^)^26^.

**Figure 2.**
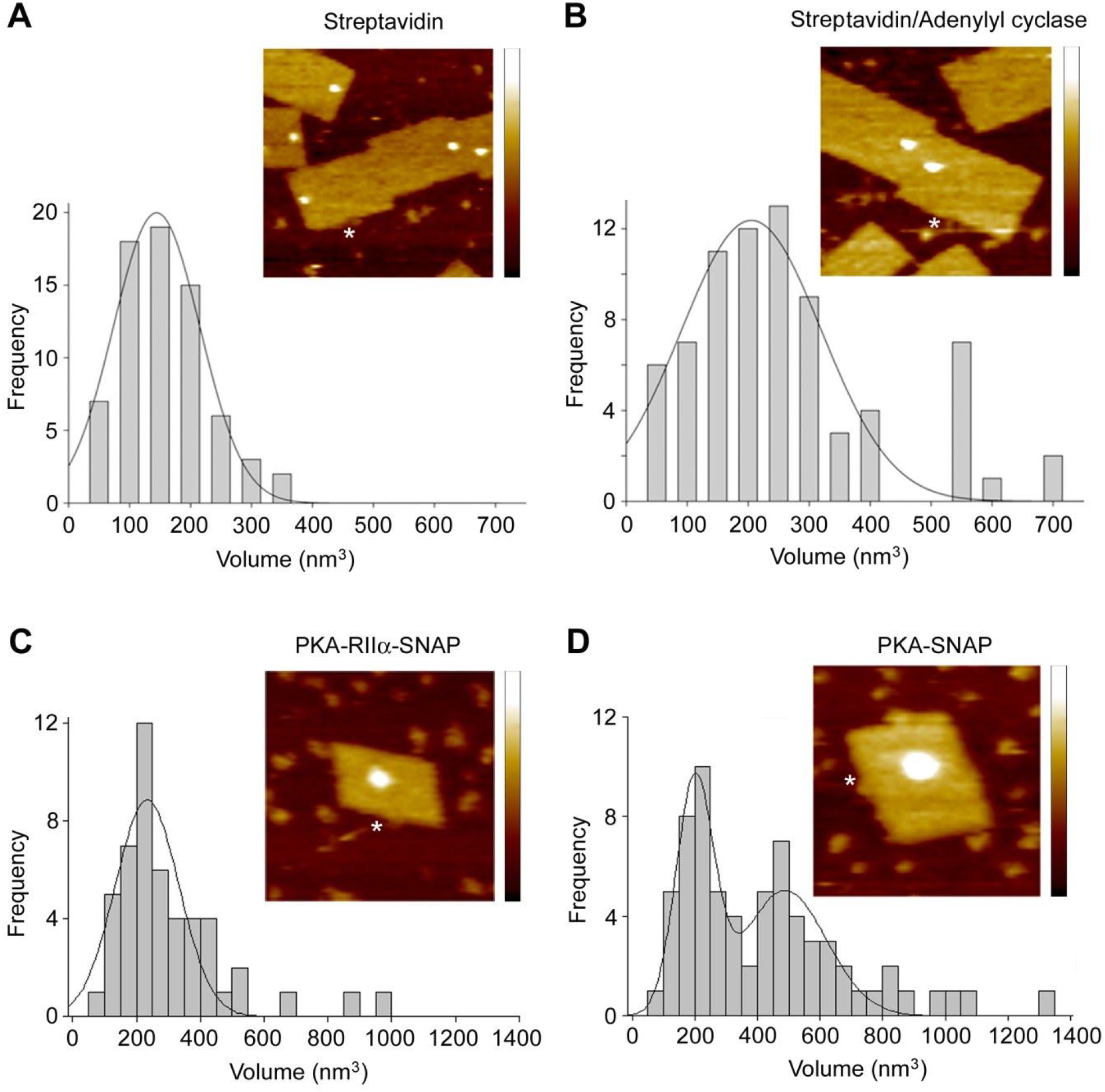
Attachment of proteins to the DNA origami tile. Insets show representative AFM images. Images are either 200 × 200 nm (**2.A-C**) or 100 × 100 nm square (**2.D**). Height range, 0-5 nm. Asterisks indicate the orientation loops. **2.A:** Volume distribution of streptavidin attached to the tile via the biotinylated staple. The peak volume corresponds to one streptavidin molecule. **2.B:** Volume distribution of attached streptavidin-bio-AC-CaM complexes. The volume increase corresponds to one bio-AC-CaM complex attached to each streptavidin molecule. **2.C:** Volume distribution of attached SNAP-tagged PKA R subunit. The peak volume corresponds to one R subunit. **2.D:** Volume distribution of attached particles after incubation of tiles with the PKA holoenzyme. The distribution shows two peaks, one corresponding to the R subunit dimer (as in **2.C**) and one to the holoenzyme.

We then incubated streptavidin-decorated tiles with a biotinylated adenylyl cyclase (bio-AC) that is derived from the catalytic domain of the CyaA toxin from *Bordetella pertussis*^27^ (Figure 2.B). To ensure the activity of the catalytic domain of bio-AC, we incubated the protein with calmodulin (CaM), prior to incubation with streptavidin. The molecular mass of the recombinant calmodulin^27^ is 19.5 kDa, and therefore the molecular mass of the bio-AC-CaM complex is 75 kDa in total. This corresponds to a theoretical volume of 142 nm^3^. The majority (68±9%; n=75) of the structures showed an increase in volume after incubation with adenylyl cyclase The average volume measured for the streptavidin-bio-AC-CaM complex (combined molecular mass,130 kDa; theoretical volume, 247 nm^3^) was 263±12 nm^3^ (n=50). The extent of the increase indicates that on average one bio-AC-CaM complex bound to each streptavidin molecule; however, the increased width of the volume distribution for this sample suggests that some streptavidin molecules might carry more than one adenylyl cyclase molecule, consistent with the fact that bound streptavidin has three sites available for binding biotin. It should be noted that as the bio-AC was biotinylated *in vivo* (see Supplementary Information), it contains a single biotin tag per enzyme molecule. All the proteins prepared and used in these experiments were characterized by SDS-polyacrylamide gel electrophoresis (SDS-PAGE; Supplementary Figure 2).

Next, R subunits of PKA were attached to the tiles in the designated position, ~20 nm away from the binding site for the streptavidin/adenylyl cyclase complex, and immediately to the left of the orientation loop. The tiles were decorated with the RIIα subunit with an efficiency of 63±16% (n=55) (Figure 2.C), and the mean volume of the attached particles was 233±11 nm^3^ (n=50). The theoretical volume of the R subunit of PKA is 123 nm^3^ for a monomer and 246 nm^3^ for a dimer, indicating that the majority of the attached particles are RIIα dimers. When the tiles were incubated with the PKA holoenzyme, the volume distribution showed two peaks, one at 200±8 nm^3^ (n=69) and another at 488±23 nm^3^ (n=68), indicating that not all the R subunit dimers were associated with C subunits to form holoenzymes; hence, a mixed population of holoenzymes and R subunits was present (Figure 2.D). The attachment efficiency of the holoenzyme on the tiles was 60±9% (n=150). The correct placement of the RIIα subunits on the tiles was confirmed by measuring the distance of the bound particles from the ends and the sides of the tiles. The observed distances were as expected given the known position of the benzylguanine-modified staple (Supplementary Figure 3). In contrast, when the tiles were incubated with C subunits of PKA, which do not bear any modification, only a very small number of decorated tiles were seen (5±2% of all well-formed tiles; n=55). Furthermore, the observed particles were in random positions on the tiles (Supplementary Figure 3). The above result confirms that the binding of the R subunit of PKA to the DNA origami tiles is precise and controlled.

We used FS-AFM to track the activation of PKA, attached alone to the tiles. Tiles bearing PKA holoenzymes were visualized at a scanning rate of 4 frames per minute in the presence of caged cAMP (1 mM). A rapid (<20 s) dissociation of the two C subunits occurred upon UV photolysis of the cage (Figure 3.A,B). In contrast, in the presence of the PKA inhibitor (R_p_)-cAMPS^28,29^ no dissociation of the PKA holoenzyme was seen (Figure 3.B,C). After photolysis of the caged cAMP, the volume of the largest attached particle (arrows, Figure 3.A) fell by 50±7% (n=6), consistent with the release of the two C subunits. Two smaller particles can be observed next to the large particle, consistent with the two catalytic subunits being released but staying in the close proximity of the regulatory dimer. In contrast, no change in the appearance or volume of the measured particles was observed in the presence of (R_p_)-cAMPS.

**Figure 3.**
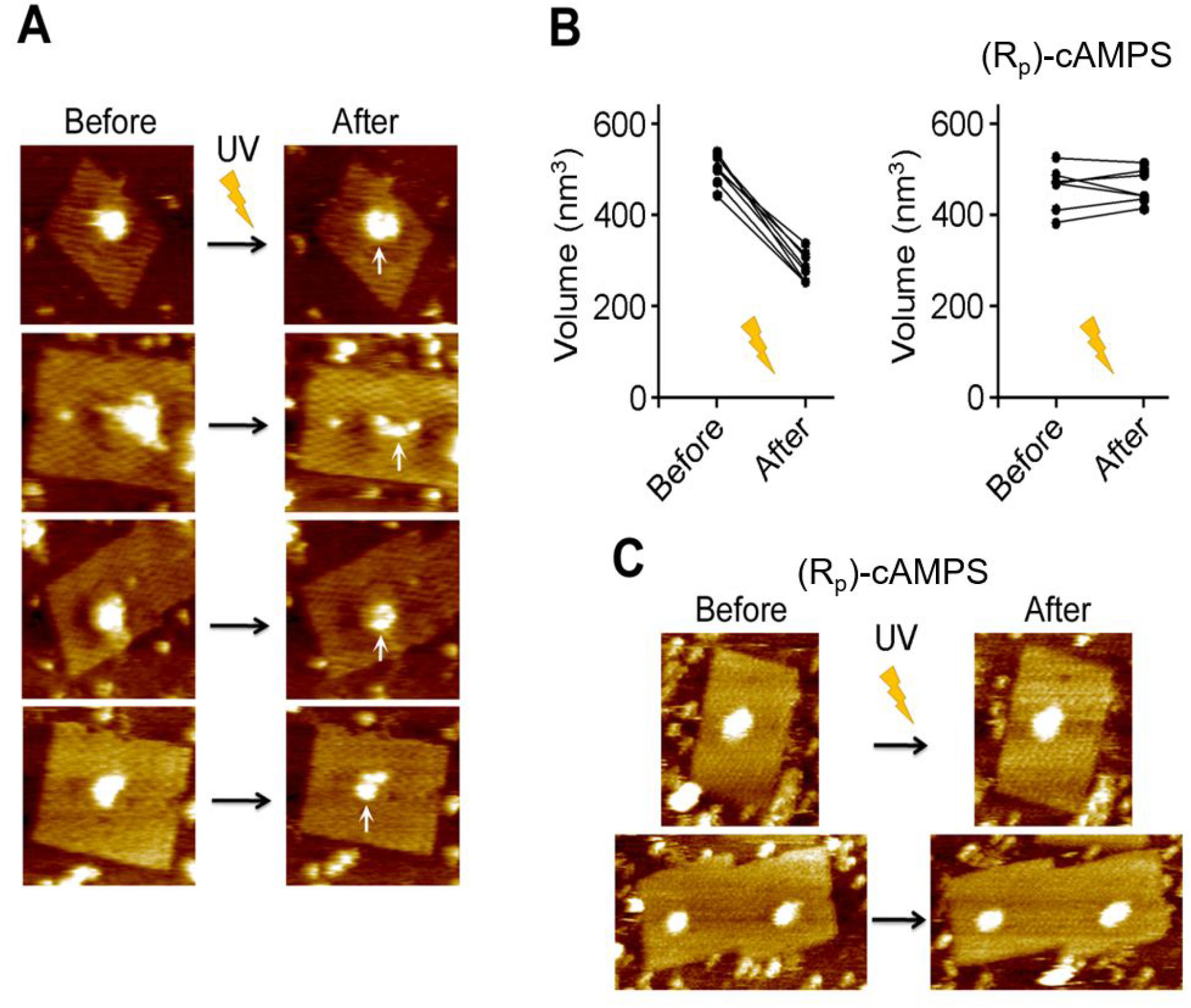
Activation of PKA by UV photolysis of caged cAMP. **3.A:** Tiles bearing a PKA holoenzyme, imaged in the presence of caged cAMP. Upon UV photolysis of the caged cAMP, indicated by the flash, the holoenzyme underwent dissociation. **3.B:** Effect of UV photolysis of caged cAMP on the volumes of PKA holoenzymes. **3.C:** Images from a control experiment in the presence of the PKA inhibitor (R_p_)-cAMPS.

We next visualized the activation of PKA by cAMP produced from ATP by adenylyl cyclase. ATP was generated by UV photolysis of caged ATP (1 mM). It was not possible in these experiments to measure the concentration of cAMP to which the bound PKA was exposed, but it is likely that it is substantially lower than that achieved by UV photolysis of caged cAMP (above). A rapid dissociation of the PKA holoenzyme was again observed (Figure 4.A,B), and the volume of the largest attached PKA particle (arrows, Figure 4.A) fell by 49±12% (n=6). The dissociation can be seen in sections taken through the attached particles (Figure 4.C). Again, no dissociation of the PKA holoenzyme was seen in the presence of (R_p_)-cAMPS (Figure 4.B,D). The distance between the central RIIα dimer and the C subunit particles in close proximity was 9.9±1.4 nm (n=13; Supplementary Figure S4), agreeing with previous studies on the dissociation of the PKA holoenzyme^21^

**Figure 4.**
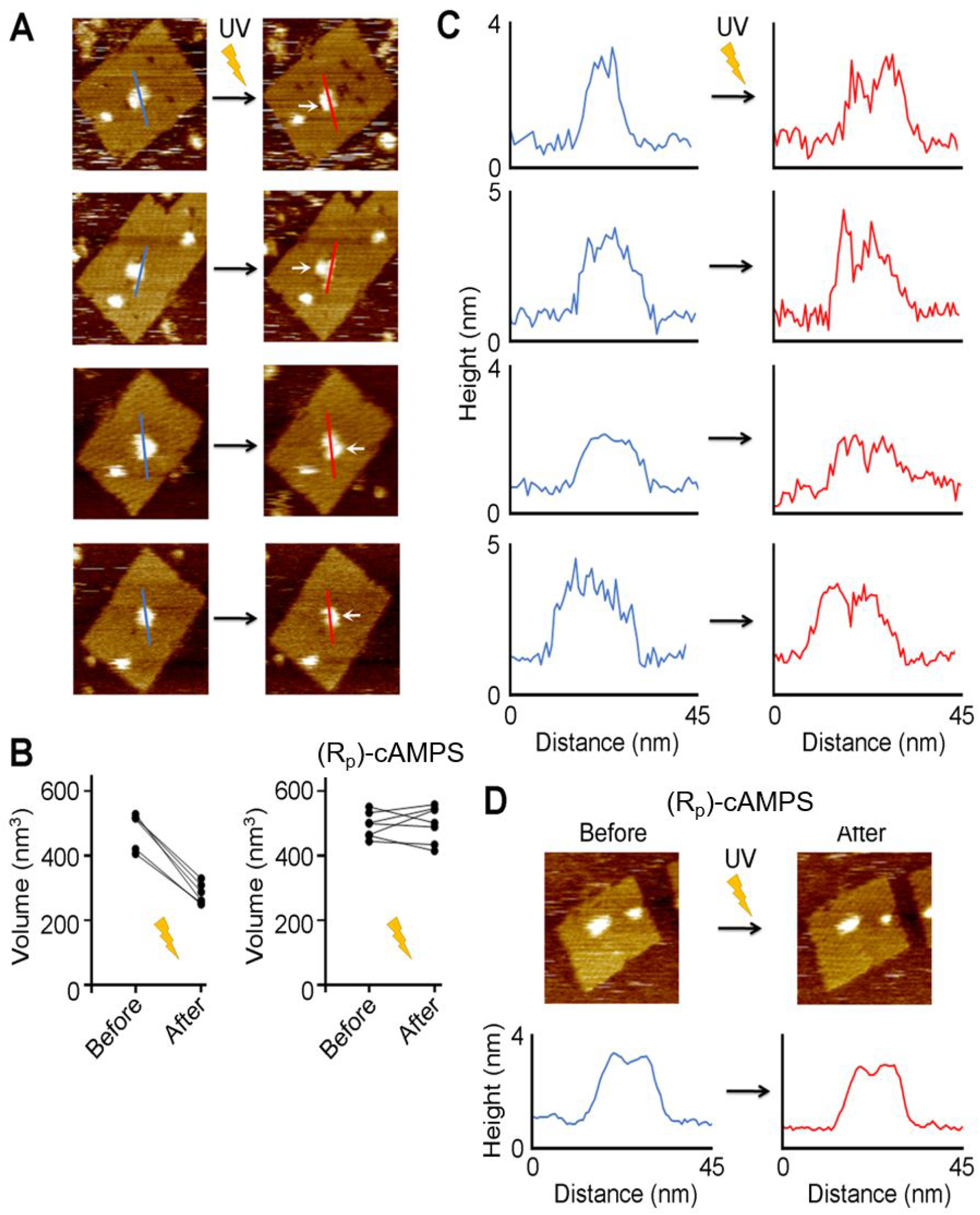
Activation of PKA using cAMP produced from caged ATP by adenylyl cyclase. **4.A:** Tiles bearing a PKA holoenzyme and the catalytic subunit of adenylyl cyclase, imaged in the presence of caged ATP. Upon UV photolysis of caged ATP, adenylyl cyclase converts ATP to cAMP and the PKA holoenzyme undergoes dissociation. **4.B:** Effect of UV photolysis of caged ATP on the volumes of PKA holoenzymes. **4.C:** Cross-sections of the holoenzyme, taken at the positions indicated by the lines in 4.a, before and after activation. **4.D:** Images from a control experiment in the presence of the PKA inhibitor (R_p_)-cAMPS.

The R and C subunits of PKA used in this study bear His_6_-tags at the C termini of the proteins. It has been shown previously that the tag does not interfere with the activity of the subunits^28^. In addition to the His_6_ tag, the R subunit was modified to bear an additional SNAP tag at the N terminus, which contains the dimerization/docking domain of the protein^30^. Nevertheless, dimerization of the RIIα subunit did not seem to be inhibited in the present study, as suggested by the volume measurements of the RIIα subunit. cAMP-induced activation of PKA has been studied before, for example using fluorescence polarization assays^31^ and hydrogen-deuterium exchange studies^32^. Further, the PKA holoenzyme has been visualized by negative-stain electron microscopy^21^. However, the effect of cAMP on PKA structure has never previously been observed at the single-molecule level. In our study, we provide a platform on which to study protein-protein interactions and parts of signalling pathways at the single-molecule level. Using this platform, we saw a clear dissociation of the PKA holoenzyme, both when PKA was activated by caged cAMP, and when activated by cAMP generated by adenylyl cyclase from ATP. The two dissociated C subunits remained on the origami tile, within ~10 nm of the R subunit dimer. Note that, as mentioned above, individual C subunits do not bind efficiently to the tiles, so their retention likely involves a continuing interaction with the R subunit dimer, as proposed by Smith *et al*.^21^.

Overall, we show that the use of fast-scan AFM to observe the behavior of proteins anchored to DNA origami tiles offers a powerful new method with which to interrogate protein function and protein-protein interactions and highlight the potential of DNA nanotechnology as a tool in the observation of signalling pathways at the single-molecule level.

## ASSOCIATED CONTENT

Direct Visualization of Protein Kinase A Activation on DNA Origami using Fast-Scan Atomic Force Microscopy.

**Supporting Information** (PDF)

## AUTHOR INFORMATION

### Author Contributions

IM designed and performed the experiments, analyzed the data and wrote the manuscript. DL, HS and ME provided resources and contributed to the experimental design and data interpretation. JME and RMH conceived the project idea, designed experiments and revised the manuscript. All authors have given approval to the final version of the manuscript.

## ACKNOWLEDGMENT

We thank Professor Susan Taylor for the kind gift of the RIIα and catalytic PKA subunit plasmids and Professor Takashii Morii for helpful discussions. HS acknowledges funding from JSPS KAKENHI (Grant Number JP16H06356). ME acknowledges funding from JSPS KAKENHI (Grant Number 18KK0139). RMH acknowledges funding from the Biotechnology and Biological Sciences Research Council Grant BB/J018236/1.

## ABBREVIATIONS

PKA: 
FS-AFM: 
SDS-PAGE: 

## Notes

### Competing Interest Statement

The authors have declared no competing interest.

## REFERENCES

(1) Yan, H. DNA-Templated Self-Assembly of Protein Arrays and Highly Conductive Nanowires. Science 2003, 301 (5641), 1882–1884. https://doi.org/10.1126/science.1089389.

(2) Chhabra, R.; Sharma, J.; Ke, Y.; Liu, Y.; Rinker, S.; Lindsay, S.; Yan, H. Spatially Addressable Multiprotein Nanoarrays Templated by Aptamer-Tagged DNA Nanoarchitectures. J. Am. Chem. Soc. 2007, 129 (34), 10304–10305. https://doi.org/10.1021/ja072410u.

(3) Williams, B. A. R.; Lund, K.; Liu, Y.; Yan, H.; Chaput, J. C. Self-Assembled Peptide Nanoarrays: An Approach to Studying Protein-Protein Interactions. Angew. Chem. Int. Ed. Engl. 2007, 46 (17), 3051–3054. https://doi.org/10.1002/anie.200603919.

(4) Ke, Y.; Lindsay, S.; Chang, Y.; Liu, Y.; Yan, H. Self-Assembled Water-Soluble Nucleic Acid Probe Tiles for Label-Free RNA Hybridization Assays. Science 2008, 319 (5860), 180–183. https://doi.org/10.1126/science.1150082.

(5) Saccà, B.; Meyer, R.; Erkelenz, M.; Kiko, K.; Arndt, A.; Schroeder, H.; Rabe, K. S.; Niemeyer, C. M. Orthogonal Protein Decoration of DNA Origami. Angew. Chem. Int. Ed. Engl. 2010, 49 (49), 9378–9383. https://doi.org/10.1002/anie.201005931.

(6) Goodman, R. P.; Erben, C. M.; Malo, J.; Ho, W. M.; McKee, M. L.; Kapanidis, A. N.; Turberfield, A. J. A Facile Method for Reversibly Linking a Recombinant Protein to DNA. Chembiochem 2009, 10 (9), 1551–1557. https://doi.org/10.1002/cbic.200900165.

(7) Fu, J.; Liu, M.; Liu, Y.; Woodbury, N. W.; Yan, H. Interenzyme Substrate Diffusion for an Enzyme Cascade Organized on Spatially Addressable DNA Nanostructures. J. Am. Chem. Soc. 2012, 134 (12), 5516–5519. https://doi.org/10.1021/ja300897h.

(8) Nakata, E.; Liew, F. F.; Uwatoko, C.; Kiyonaka, S.; Mori, Y.; Katsuda, Y.; Endo, M.; Sugiyama, H.; Morii, T. Zinc-Finger Proteins for Site-Specific Protein Positioning on DNA-Origami Structures. Angew. Chemie Int. Ed. 2012, 51 (10), 2421–2424. https://doi.org/10.1002/anie.201108199.

(9) Wilner, O. I.; Shimron, S.; Weizmann, Y.; Wang, Z.-G.; Willner, I. Self-Assembly of Enzymes on DNA Scaffolds: En Route to Biocatalytic Cascades and the Synthesis of Metallic Nanowires. Nano Lett. 2009, 9 (5), 2040–2043. https://doi.org/10.1021/nl900302z.

(10) Endo, M.; Katsuda, Y.; Hidaka, K.; Sugiyama, H. Regulation of DNA Methylation Using Different Tensions of Double Strands Constructed in a Defined DNA Nanostructure. J. Am. Chem. Soc. 2010, 132 (5), 1592–1597. https://doi.org/10.1021/ja907649w.

(11) Derr, N. D.; Goodman, B. S.; Jungmann, R.; Leschziner, A. E.; Shih, W. M.; Reck-Peterson, S. L. Tug-of-War in Motor Protein Ensembles Revealed with a Programmable DNA Origami Scaffold. Science 2012, 338 (6107), 662–665. https://doi.org/10.1126/science.1226734.

(12) Walsh, D. A.; Perkins, J. P.; Krebs, E. G. An Adenosine 3’,5’-Monophosphate-Dependant Protein Kinase from Rabbit Skeletal Muscle. J. Biol. Chem. 1968, 243 (13), 3763–3765.

(13) Blume-Jensen, P.; Hunter, T. Oncogenic Kinase Signalling. Nature 2001, 411 (6835), 355–365. https://doi.org/10.1038/35077225.

(14) Enns, L. C.; Ladiges, W. Protein Kinase A Signaling as an Anti-Aging Target. Ageing Res. Rev. 2010, 9 (3), 269–272. https://doi.org/10.1016/j.arr.2010.02.004.

(15) Gill, G. N.; Garren, L. D. Role of the Receptor in the Mechanism of Action of Adenosine 3’:5’-Cyclic Monophosphate. Proc. Natl. Acad. Sci. 1971, 68 (4), 786–790. https://doi.org/10.1073/pnas.68.4.786.

(16) Rubin, C. S. A Kinase Anchor Proteins and the Intracellular Targeting of Signals Carried by Cyclic AMP. Biochim. Biophys. Acta 1994, 1224 (3), 467–479.

(17) Herberg, F. W.; Taylor, S. S.; Dostmann, W. R. Active Site Mutations Define the Pathway for the Cooperative Activation of CAMP-Dependent Protein Kinase. Biochemistry 1996, 35 (9), 2934–2942. https://doi.org/10.1021/bi951647c.

(18) Skålhegg, B. S.; Taskén, K. Specificity in the CAMP/PKA Signaling Pathway. Differential Expression, Regulation, and Subcellular Localization of Subunits of PKA. Front. Biosci. 1997, 2, d331–42.

(19) Colledge, M.; Scott, J. D. AKAPs: From Structure to Function. Trends Cell Biol. 1999, 9 (6), 216–221.

(20) Taylor, S. S.; Buechler, J. A.; Yonemoto, W. CAMP-Dependent Protein Kinase: Framework for a Diverse Family of Regulatory Enzymes. Annu. Rev. Biochem. 1990, 59 (1), 971–1005. https://doi.org/10.1146/annurev.bi.59.070190.004543.

(21) Smith, F. D.; Esseltine, J. L.; Nygren, P. J.; Veesler, D.; Byrne, D. P.; Vonderach, M.; Strashnov, I.; Eyers, C. E.; Eyers, P. A.; Langeberg, L. K.; Scott, J. D. Local Protein Kinase A Action Proceeds through Intact Holoenzymes. Science 2017, 356 (6344), 1288–1293. https://doi.org/10.1126/science.aaj1669.

(22) Walker-Gray, R.; Stengel, F.; Gold, M. G. Mechanisms for Restraining CAMP-Dependent Protein Kinase Revealed by Subunit Quantitation and Cross-Linking Approaches. Proc. Natl. Acad. Sci. U. S. A. 2017, 114 (39), 10414–10419. https://doi.org/10.1073/pnas.1701782114.

(23) Green, N.; Avidin. 3. The nature of the biotin-binding site. Biochem. J. 1963, 89 (3), 599–609. https://doi.org/10.1042/bj0890599.

(24) Kuzuya, A.; Komiyama, M. Design and Construction of a Box-Shaped 3D-DNA Origami. Chem. Commun. 2009, No. 28, 4182. https://doi.org/10.1039/b907800b.

(25) Voigt, N. V; Tørring, T.; Rotaru, A.; Jacobsen, M. F.; Ravnsbaek, J. B.; Subramani, R.; Mamdouh, W.; Kjems, J.; Mokhir, A.; Besenbacher, F.; Gothelf, K. V. Single-Molecule Chemical Reactions on DNA Origami. Nat. Nanotechnol. 2010, 5 (3), 200–203. https://doi.org/10.1038/nnano.2010.5.

(26) Schneider, S. W.; Lärmer, J.; Henderson, R. M.; Oberleithner, H. Molecular Weights of Individual Proteins Correlate with Molecular Volumes Measured by Atomic Force Microscopy. Pflugers Arch. Eur. J. Physiol. 1998. https://doi.org/10.1007/s004240050524.

(27) Selwa, E.; Davi, M.; Chenal, A.; Sotomayor-Pérez, A. C.; Ladant, D.; Malliavin, T. E. Allosteric Activation of *Bordetella Pertussis* Adenylyl Cyclase by Calmodulin: Molecular Dynamics and Mutagenesis Studies. J. Biol. Chem. 2014, 289 (30), 21131–21141.https://doi.org/10.1074/jbc.M113.530410.

(28) Dostmann, W. R. G. (RP)-CAMPS Inhibits the CAMP-Dependent Protein Kinase by Blocking the cAMP-Induced Conformational Transition. FEBS Lett. 1995, 375 (3), 231–234. https://doi.org/10.1016/0014-5793(95)01201-O.

(29) Wu, J.; Jones, J. M.; Xuong, N. H.; Ten Eyck, L. F.; Taylor, S. S. Crystal Structures of RIα Subunit of Cyclic Adenosine 5’-Monophosphate (CAMP)-Dependent Protein Kinase Complexed with (R p)-Adenosine 3’,5’-Cyclic Monophosphothioate and (S p)-Adenosine 3’,5’-Cyclic Monophosphothioate, the Phosphothioate Analogues of CAMP. Biochemistry 2004, 43 (21), 6620–6629. https://doi.org/10.1021/bi0302503.

(30) Narayana, N.; Cox, S.; Shaltiel, S.; Taylor, S. S.; Xuong, N. Crystal Structure of a Polyhistidine-Tagged Recombinant Catalytic Subunit of CAMP-Dependent Protein Kinase Complexed with the Peptide Inhibitor PKI(5-24) and Adenosine. Biochemistry 1997, 36 (15), 4438–4448. https://doi.org/10.1021/bi961947+.

(31) Banky, P.; Huang, L. J.; Taylor, S. S. Dimerization/Docking Domain of the Type Ialpha Regulatory Subunit of CAMP-Dependent Protein Kinase. Requirements for Dimerization and Docking Are Distinct but Overlapping. J. Biol. Chem. 1998, 273 (52), 35048–35055. https://doi.org/10.1074/jbc.273.52.35048.

(32) Kim, C.; Cheng, C. Y.; Saldanha, S. A.; Taylor, S. S. PKA-I Holoenzyme Structure Reveals a Mechanism for CAMP-Dependent Activation. Cell 2007, 130 (6), 1032–1043. https://doi.org/10.1016/j.cell.2007.07.018.

