## Supplementary Data for "Direct Visualization of Protein Kinase A Activation on DNA Origami using Fast-Scan Atomic Force Microscopy"

### Supporting information

#### Table of contents

**Section 1. Synthesis and characterization of DNA origami nanostructures**

**Section 2. Protein preparation**

**Section 3. Protein attachment onto DNA origami nanostructures**

**Section 4. AFM imaging and data analysis**

#### Section 1. Synthesis of DNA origami nanostructures

##### 1.1 Synthesis of DNA origami nanostructures

The DNA origami tiles (Figure S1) were designed as described in by Rothemund[1]. All staple oligonucleotides were purchased from IDT. The sample solution (20 µl) containing M13mp18 single-stranded DNA (New England Biolabs, 0.01 µM), staple DNA strands (5 equiv, 0.05 µM), Tris-HCl (pH 7.6, 20 mM), EDTA (1 mM), and $MgCl_2$ (10 mM) was annealed from 85 to 25°C at a rate of -1°C/ min, by using a thermal cycler. After annealing, excess oligonucleotides were removed using a Micro Biospin column (Bio-Rad) packed with Sephacryl S-300 (GE Healthcare).

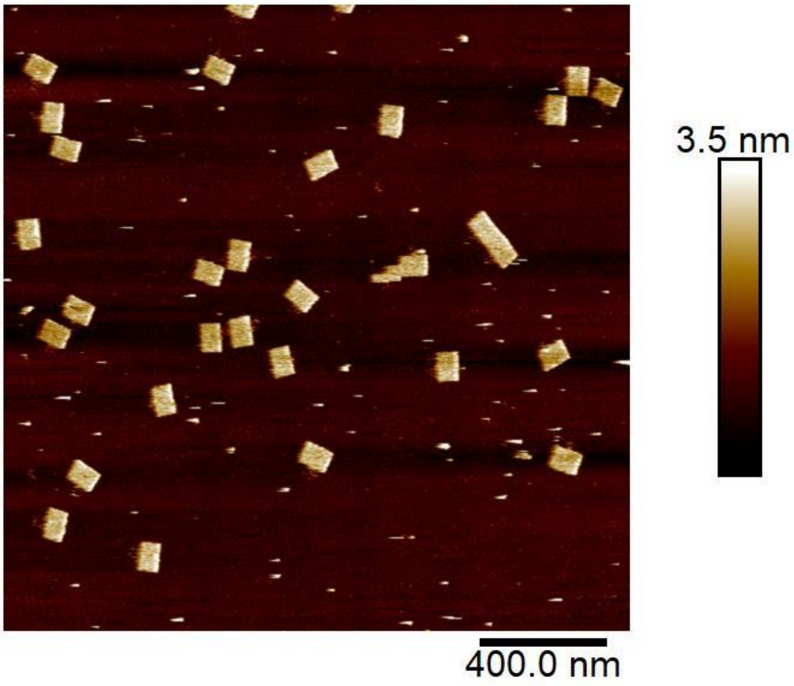


**Figure S1:** Large field of view (2 x 2 µm) of DNA origami nanostructures. The nanostructures assemble with an efficiency of 92±5%.

##### 1.2 Modifications to the DNA origami tiles

Two oligonucleotides were modified to carry one biotin and one benzylguanine modification respectively, positioned ~20 nm apart.

The sequence of the biotin functionalized ologonucleotide was:

x-5’-TTTTTTTCAAAATTAAAGTACGGTGTCTGGAAGAGGTCA-3’,

where x is the position of the biotin attachment.

The benzylguanine oligonucleotide was made using an amine functionalized oligonucleotide, as described below, and the sequence was:

5’-GCCAGACCATTACCAGTTTGATTTTCTTAACTTTTTTTT-3’-y,

where y, is the amine modification.

##### 1.3 Preparation of the benzylguanine staple

Amine-modified staples were reacted with BG-GLA-NHS (New England Biolabs). BG-GLA-NHS was diluted in DMF to a final concentration of 5 mM. 20 μl of the amine-functionalized DNA oligonucleotide (50 μM) was mixed with 10 μl of BG-GLA-NHS, 5 μl of Tris-HCl, pH 8 (1 M) and 10 μl DMF. The volume of the solution was made up to 50 μl using 5 μl of MilliQ water and the mixture was incubated overnight at 30°C. The reacted staples were purified using HPLC and then evaporated for 1 h and stored at -80°C. Subsequently, the samples were freeze-dried overnight and then resuspended in 50 μl of origami buffer.

#### Section 2. Protein preparation and purification

##### 2.1 PKA Purification and holoenzyme formation

The RIIα subunit was modified to carry a SNAP tag, by sub-cloning the fragment of the plasmid encoding the RIIα subunit of PKA into a pSNAP-tag (T7)-2 vector (New England Biolabs). The RIIα and catalytic subunits of PKA were purified as described previously[2]. In the case of RIIa, a cAMP resin was used, in order to remove residual cAMP bound to the protein, as described in Wu *et al.*[3]. The R and C subunits were mixed in a ratio of 1:1.2 in regenerated cellulose SnakeSkin dialysis membrane (Thermo Scientific: molecular mass cut-off of 10 kDa) and then dialysed overnight against 2 l of buffer containing 10 mM MOPS (pH 7), 50 mM NaCl and 1 mM DTT, with two buffer changes. The dialysed sample was collected and stored on ice at 4°C.

**2.2 Biotinylated adenylyl cyclase purification.**

The recombinant biotinylated adenylyl cyclase (bio-AC) was produced in *Escherichia coli* from plasmid pTRAC-Biotin that codes for the first 384 codons of *Bordetella pertussis* CyaA toxin fused to the *E. coli* biotin carboxyl carrier protein (BCCP). Plasmid pTRAC-Biotin was constructed by inserting the BCCP coding region - amplified from plasmid PinPoint$^{TM}$ Xa-1 (Promega Corporation, USA) - into the NdeI site from vector pTRAC-384GK[4]. Plasmid pTRAC-Biotin was co-transformed with plasmid pBirA-Cm (Avidity, LLC**,** USA; codes for the *E. coli* biotin holoenzyme synthetase, BirA) into *E. coli* BLR competent cells. Recombinant clones were grown at 30°C to mid-log phase in LB supplemented with ampicillin and chloramphenicol. Then biotin (50 μM) and IPTG (0.2 mM ; to trigger expression of BirA) were added and the culture was continued at 37°C (to trigger expression of Bccp-AC fusion) for 4 h. Collected cells were lysed by sonication and the recombinant bioAC protein was purified to homogeneity from inclusion bodies as described previously[4]. Biotinylation was confirmed by protein dot-blots and ELISA assays. Catalytic activity was measured as described in Selwa *et al.*[5]. The purified recombinant bioAC protein (theoretical molecular mass, 55.6 kDa) exhibited a $k_{cat}$ ranging from 2,000 to 3,000 $s^{-1}$ when assayed in the presence of 1 μM calmodulin. (Additional details on plasmid construction and protein production can be provided upon request).

**2.3 Characterization of proteins and protein-oligonucleotide attachment**

We characterized the four proteins used in the present study by SDS-PAGE: streptavidin, adenylyl cyclase, the RIIα subunit of PKA, modified to carry a SNAP tag, and the C subunit of PKA (Figure S2.A), were found to run at the expected molecular masses (note that streptavidin dissociates into its four subunits). We also verified the ability of a benzylguanine-modified DNA staple (32 bp) to attach to the R subunit of PKA (Figure S2.B, red rectangle). Benzylguanine forms a covalent bond with the SNAP tag on the RIIα subunit and is therefore detectable by SDS-PAGE.

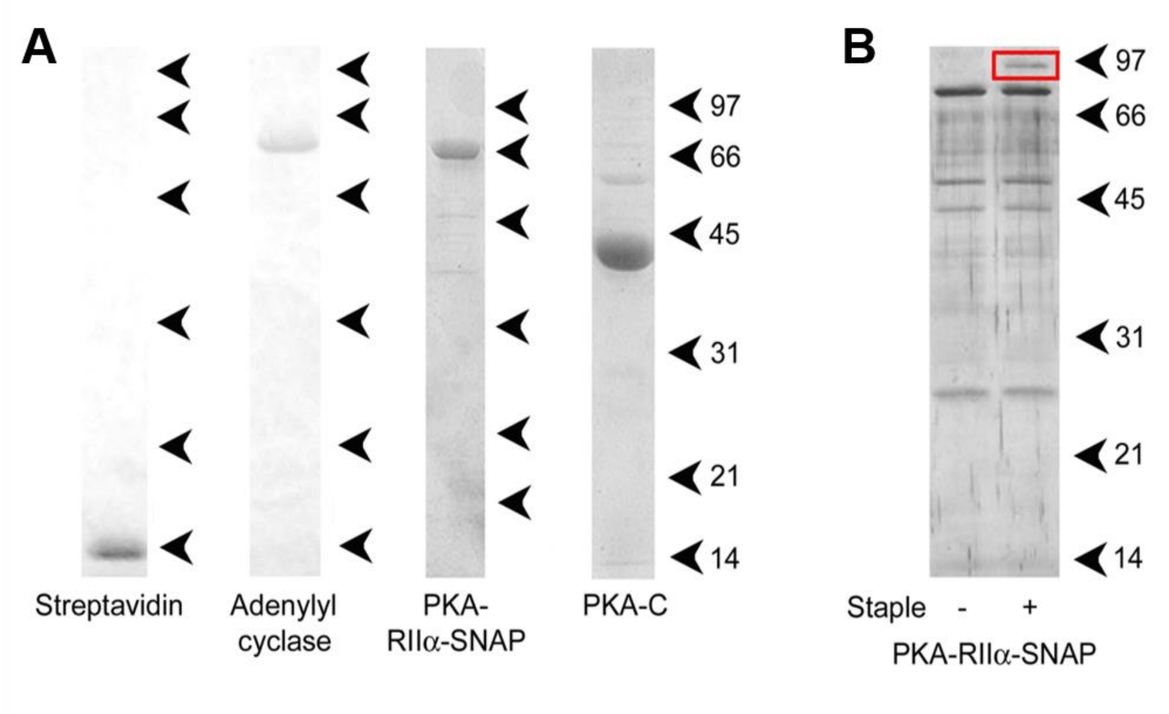


**Figure S2:** SDS-PAGE analysis of the proteins used in this study. **S2.A:** Streptavidin (with the four subunits dissociated), bio-AC, the RIIα subunit of PKA, modified to carry a SNAP tag, and the C subunit of PKA. **S2.B:** Confirmation of the successful binding of a benzylguanine-modified oligonuclotide to the regulatory subunit of PKA. In the sample incubated with the oligo, an extra band can be seen, corresponding to a 32 bp oligonucleotide attached to the protein.

#### Section 3. Protein attachment on DNA origami nanostructures

##### 3.1 Silver-staining of protein-DNA complexes

The benzylguanine-modified staple was diluted ten times in SNAP labelling buffer (50 mM Tris Cl (pH 7.5), 100 mM NaCl, 1 mM DTT), to a final concentration of 0.05 μM and mixed with purified RIIα subunit. Each sample was incubated for 2 h at room temperature. The RIIα-staple strand complexes were analysed by SDS-PAGE followed by silver staining.

##### 3.2 Adenylyl cyclase attachment to DNA origami

Assembled tiles were incubated with 0.17 μM streptavidin (Sigma-Aldrich) for 5 min, and filtered using a Millipore filter (Millipore, MA, USA) unit with molecular mass cut-off of 100 kDa to remove free protein. Streptavidin-decorated tiles were incubated with bio-AC-CaM (0.24 μM bio-AC plus 0.5 μM CaM) for 15 min. The tiles were then filtered as above and imaged.

##### 3.3 RIIa/PKA attachment to DNA origami

Protein (either the RIIα subunit or PKA holoenzyme) was diluted in SNAP labelling buffer (50 mM Tris Cl (pH 7.5), 100 mM NaCl, 1 mM DTT), and 10 μl of the solution was mixed with 10 μl of DNA origami tiles. The sample was incubated for 2 h at room temperature and then diluted ten times in SNAP labelling buffer containing 10 mM $MgCl_2$. The sample (45 μl) was then deposited on freshly cleaved mica and was incubated for 5 min at room temperature; excess sample was washed five times with 1 ml SNAP labelling buffer (Figure S3).

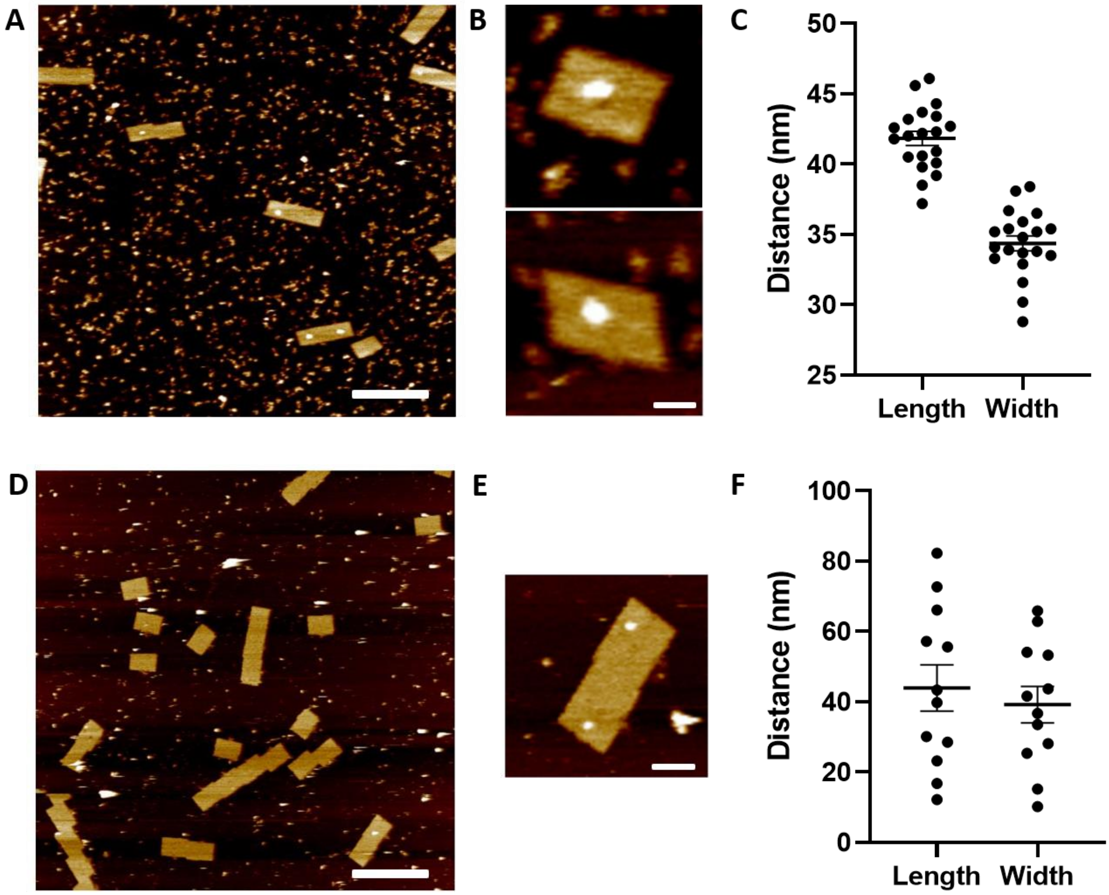


**Figure S3: Positioning of proteins on the DNA origami platform. S3.A:** Large (2 x 2 μm) field of view of DNA origami tiles decorated with the RIIα subunit of PKA. **S3.B:** High-magnification images of the tiles showing the protein attached in the expected position. **S3.C:** Measurements of the position of the protein in relation to the length and width of the tile show that the attached particles are present in the expected position of the tiles, with the small variations likely rising from the existence of a flexible linker that attaches the protein to the tile and the orientation of the protein in each image. **S3.D:** Large (2 x 2 μm) field of view of DNA origami tiles decorated with the catalytic subunit of PKA. **S3.E:** High-magnification images of the tiles showing proteins present in random positions on the tile. **S3.F:** Measurements of the position of the protein in relation to the length and width of the tile show that the attached particles are present at random on the tiles. The low number of particles present and the random positions of the particles, indicate that binding of proteins onto the DNA origami platform is driven by the existence of specific tags. Scale bars: 400 nm for a, d, 50 nm for b and 70 nm for e. Height range: 0 – 4 nm for all images.

##### 3.4 Activation of PKA

Either 1 mM caged cAMP (4,5-dimethoxy-2-nitrobenzyl-cAMP; Invitrogen) or 1 mM caged ATP (adenosine 5′-triphosphate γ-(1-[2-nitrophenyl]ethyl) ester; Molecular Probes) was added to the sample prior to imaging. The caged nucleotides were activated at a set time during imaging, using UV flash photolysis.

When PKA was activated the catalytic subunits  observed to be still present on the DNA origami tile. Measuring the distance between the regulatory dimer and the catalytic subunits showed us a consistent mean distance of 9.9±1.4 nm (n=13, Figure S4), which is in agreement with previous studies[6].

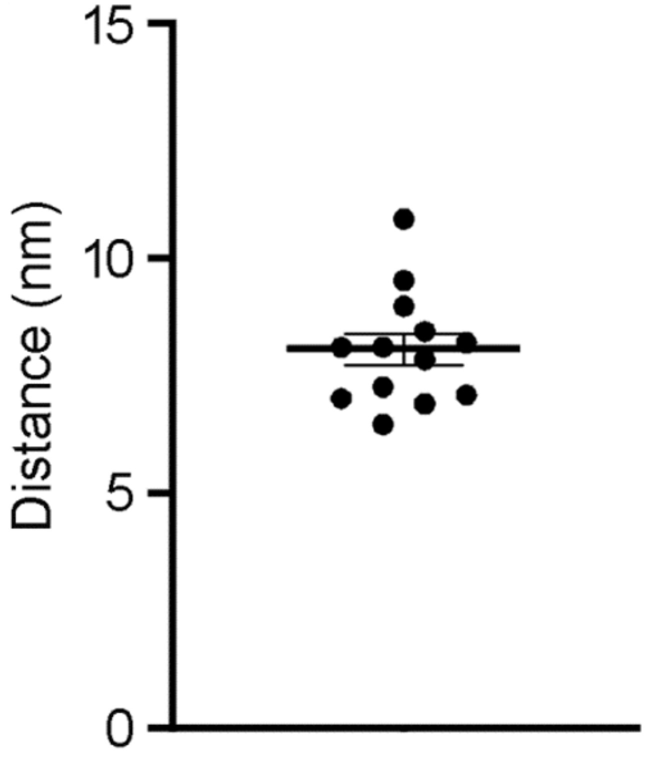


**Figure S4:** Distances between the central RIIα dimer and the C subunit particles in close proximity.

#### Section 4. AFM imaging and data analysis

##### 4.1 AFM imaging

Imaging was performed using a Dimension FastScan microscope (Bruker) equipped with a flash photolysis module, under fluid conditions. The probes used were the FastScan-D probes (Bruker), with a resonant frequency of 90 kHz, a spring constant of 0.21 $Nm^{-1}$ and a nominal tip radius of 8 nm.

##### 4.2 Data analysis

Images of all datasets were plane-fitted using the speed-optimized plane correction function of the SPIP software (Image Metrology A/S, Hørsholm, Denmark), which fits each line in the horizontal axis to a polynomial equation. SPIP was also used for calculation of the volumes of proteins attached to DNA origami tile. In this case, the 'inspection window' feature of SPIP was used to zoom into individual tiles and then the 'circular area of interest' tool was used to allow the software to calculate only the volume of the protein rather than that of the whole tile. SPIP uses the following equation for the calculation of volume:

$$Z_{net\ volume} = Z_{material\ volume} - Z_{void\ volume},$$

where $Z_{material\ volume}$ is the volume of all pixels inside the shape's contour with a Z value greater than or equal to zero:

$$Z_{material\ volume} = \sum_{\{Z(x,y) \in shape | Z \geq 0\}} Z(x,y)dxdy,$$

where dx and dy are the point spacings in the X and Y directions of the image, respectively.

$Z_{void\ volume}$ is the volume of all pixels inside the shape's contour with a Z value lower than or equal to zero:

$$Z_{\text{void volume}} = \sum_{\{Z(x,y)\in \text{shape}|Z\leq 0\}} Z(x,y)dxdy ,$$

where dx and dy are the point spacing in the x and y directions of the image, respectively.

SPIP was also used to draw cross-sections of sample features, for measurement of the dimensions of the tiles, and for the 3D rendering of the images.

Volume histograms were drawn with bin widths chosen according to Scott's equation[7]:

$$\text{Bin width} = 3.5\sigma/n^{1/3},$$

where σ is an estimate of the standard deviation and *n* is the sample size.

Where Gaussian curves were fitted to the data, the curve fitting tool of SigmaPlot (Systat Software Inc., California, USA) was used. The following equations were used by the software to fit either one or two peak curves to the histogram by iteration:

One distribution:

$$f = a_1 \exp\left(\frac{-0.5(x - x_1)}{b_1^2}\right),$$

Two distributions:

$$f = a_1 \exp\left(\frac{-0.5(x - x_1)}{b_1^2}\right) + a_2 \exp\left(\frac{-0.5(x - x_2)}{b_2^2}\right),$$

where $a_1$ and $b_1$ are the coefficients for distribution 1, $a_2$ and $b_2$ are the coefficients for distribution 2 and $x_1$ and $x_2$ are the means of the distributions.

Molecular volumes of proteins based on molecular mass were calculated using the equation proposed by Schneider *et al.*[8].

(1) Rothemund, P. W. K. Folding DNA to Create Nanoscale Shapes and Patterns. *Nature* **2006**, *440* (7082), 297–302. https://doi.org/10.1038/nature04586.

(2) Saraswat, L. D.; Filutowics, M.; Taylor, S. [31] Expression and Mutagenesis of the Regulatory Subunit of CAMP-Dependent Protein Kinase in Escherichia Coli. *Methods Enzymol.* **1988**, *159*, 325–336. https://doi.org/10.1016/0076-6879(88)59033-X.

(3) Wu, J.; Brown, S.; Xuong, N.-H.; Taylor, S. S. RIα Subunit of PKA: A CAMP-Free Structure Reveals a Hydrophobic Capping Mechanism for Docking CAMP into Site B. *Structure* **2004**, *12* (6), 1057–1065. https://doi.org/10.1016/J.STR.2004.03.022.

(4) Vougier, S.; Mary, J.; Dautin, N.; Vinh, J.; Friguet, B.; Ladant, D. Essential Role of Methionine Residues in Calmodulin Binding to Bordetella Pertussis Adenylate Cyclase, as Probed by Selective Oxidation and Repair by the Peptide Methionine Sulfoxide Reductases. *J. Biol. Chem.* **2004**, *279* (29), 30210–30218. https://doi.org/10.1074/jbc.M400604200.

(5) Selwa, E.; Davi, M.; Chenal, A.; Sotomayor-Pérez, A. C.; Ladant, D.; Malliavin, T. E. Allosteric Activation of *Bordetella Pertussis* Adenylyl Cyclase by Calmodulin: Molecular Dynamics and Mutagenesis Studies. *J. Biol. Chem.* **2014**, *289* (30), 21131–21141. https://doi.org/10.1074/jbc.M113.530410.

(6) Smith, F. D.; Esseltine, J. L.; Nygren, P. J.; Veesler, D.; Byrne, D. P.; Vonderach, M.; Strashnov, I.; Eyers, C. E.; Eyers, P. A.; Langeberg, L. K.; Scott, J. D. Local Protein Kinase A Action Proceeds through Intact Holoenzymes. *Science (80-. ).* **2017**, *356* (6344), 1288–1293. https://doi.org/10.1126/science.aaj1669.

(7) SCOTT, D. W. On Optimal and Data-Based Histograms. *Biometrika* **1979**, *66* (3), 605–610. https://doi.org/10.1093/biomet/66.3.605.

(8) Schneider, S. W.; Lärmer, J.; Henderson, R. M.; Oberleithner, H. Molecular Weights of Individual Proteins Correlate with Molecular Volumes Measured by Atomic Force Microscopy. *Pflugers Arch. Eur. J. Physiol.* **1998**. https://doi.org/10.1007/s004240050524.
